# Vasoactive Intestinal Peptide Amphiphile Micelle Chemical Structure and Hydrophobic Domain Influence Immunomodulatory Potentiation

**DOI:** 10.1101/2021.09.10.459855

**Authors:** Xiaofei Wang, Rui Zhang, Bryce D. Lindaman, Caitlin N. Leeper, Adam G. Schrum, Bret D. Ulery

**Affiliations:** Department of Biomedical, Biological, and Chemical Engineering, University of Missouri, Columbia, MO; Department of Molecular Microbiology and Immunology, University of Missouri, Columbia, MO; Department of Surgery, University of Missouri, Columbia, MO

## Abstract

Vasoactive intestinal peptide (VIP) is a neuropeptide capable of downregulating innate immune responses in antigen presenting cells (APCs) by suppressing their pro-inflammatory cytokine secretion and cell surface marker expression. Though VIP’s bioactivity could possibly be leveraged as a treatment for autoimmune disorders and transplant tolerance, drug delivery innovation is required to overcome its intrinsically limited cellular delivery capacity due to its short *in vivo* lifetime. One option is to employ peptide amphiphiles (PAs) which are lipidated peptides capable of self-assembling into micelles in water that can enhance cellular association. With this approach in mind, a series of triblock VIP amphiphiles (VIPAs) has been synthesized to explore the influence of block arrangement and hydrophobicity on micelle biocompatibility and bioactivity. VIPA formulation has been found to influence the shape, size, and surface charge of VIPA micelles (VIPAMs) as well as their cytotoxicity and immunomodulatory effects. Specifically, the enclosed work provides strong evidence that cylindrical VIPAMs with aspect ratios of 1.5 - 150 and moderate positive surface charge are able to potentiate the bioactivity of VIP limiting TNF-α secretion and MHC II and CD86 surface expression on APCs. With this criteria, we have identified PalmK-(EK)_4_-VIP as our lead formulation, which showed comparable or enhanced anti-inflammatory effects relative to the unmodified VIP at all dosages evaluated. Additionally, the relationships between peptide block location and lipid block size provide further information on the chemistry-structure-function relationships of peptide amphiphile micelles for the delivery of VIP as well as potentially for other peptides more broadly.

## Introduction

Vasoactive intestinal peptide (VIP) is an exciting biologic capable of modulating innate immune responses by altering the cytokines released from antigen presenting cells (APC). Research has shown that VIP is able to specifically downregulate pro-inflammatory cytokines that are secreted from macrophages (Mφs) and dendritic cells (DCs) during their activation like tumor necrosis factor alpha (TNF-α) [1, 2]. Based on the desirable anti-inflammatory effect of VIP, considerable research has focused on utilizing this peptide for the treatment of a variety of immune pathology-related diseases including sarcoidosis [2, 3], multiple sclerosis [4–6], arthritis [7–9], and asthma [10–12]. Though functionally promising, VIP has a short half-life and a limited retention time in plasma [13] leading to low efficiency on-target peptide delivery, highly limiting its biomedical utility. Thus, an effective delivery strategy must be designed to enhance the clinical translatability of VIP.

Peptide amphiphiles (PAs) are lipidated peptides with unique amphiphilicity that allows for their self-assembly into various architectures. In aqueous solutions, the hydrophilic peptide tends to disperse in solution, but the hydrophobic lipid tends to sequester itself. When the PA is in an abundant concentration, enough hydrophobic interactions across PAs are present to facilitate lipid clustering, for which the hydrophilic peptides shield the lipids from the water overcoming the organizational entropic penalty facilitating their self-assembly into peptide amphiphile micelles (PAMs) [14]. Moreover, various lipid structures can stabilize the peptide [15, 16] and, due to the similarity of the hydrophobic lipid to the phospholipid membrane of cells, PAs can also facilitate enhanced interactions of their peptides with the cell membrane improving peptide delivery efficiency [17]. The unique structure and properties of PAs make them broadly applicable for a variety of biomedical applications including tumor detection [18], anti-cancer therapy [19, 20], and vaccine design [21–23]. While most early research in the field has focused on the use of diblock PAs comprised of just lipids and bioactive peptides, recent research has uncovered that triblock PAs containing an additional functional block (*e.g.*, hydrogen bond enabling) can enhance micellization and the formation of structures not previously achievable with diblock PAs [21, 24]. Excitingly, our recent research has shown that electrostatically complexed triblock PAM structures can greatly influence the immunogenicity of incorporated vaccine antigen peptides in a size and structure dependent manner [21, 22].

Building on this discovery, we have investigated the capacity of PAMs to enhance the bioactivity of VIP [25]. This initial work provided evidence that triblock VIP amphiphiles (VIPAs) containing a zwitterion-like block assemble into braided cylindrical VIP amphiphile micelles (VIPAMs) which suppress the pro-inflammatory behavior of mature Mφs and DCs [26]. While promising, further research is needed to investigate the influence varying lipid block hydrophobicity and PA zwitterion block arrangement have on VIPAM structure and bioactivity. In this study, four VIPAs (*i.e.*, one or two palmitoyl tails with internal or external zwitterion-like peptide block location) were chemically synthesized, physically assessed, and biologically evaluated to determine the impact micelle core hydrophobicity and size/shape have on potentiating VIP bioactivity.

## Results and Discussion

### Chemical structure and physical properties of VIPAs

The four triblock VIPAs investigated include products with a single lipid, internal zwitterion-like peptide block (PalmK-(EK)_4_-VIP), a single lipid, external zwitterion-like peptide block (PalmK-VIP-(KE)_4_), a double lipid, internal zwitterion-like peptide block (Palm_2_K-(EK)_4_-VIP), and a double lipid, external zwitterion-like peptide block (Palm_2_K-VIP-(KE)_4_). **Table 1** outlines the chemical structures of the four different VIPAs that were used in this work.

**Table 1.**
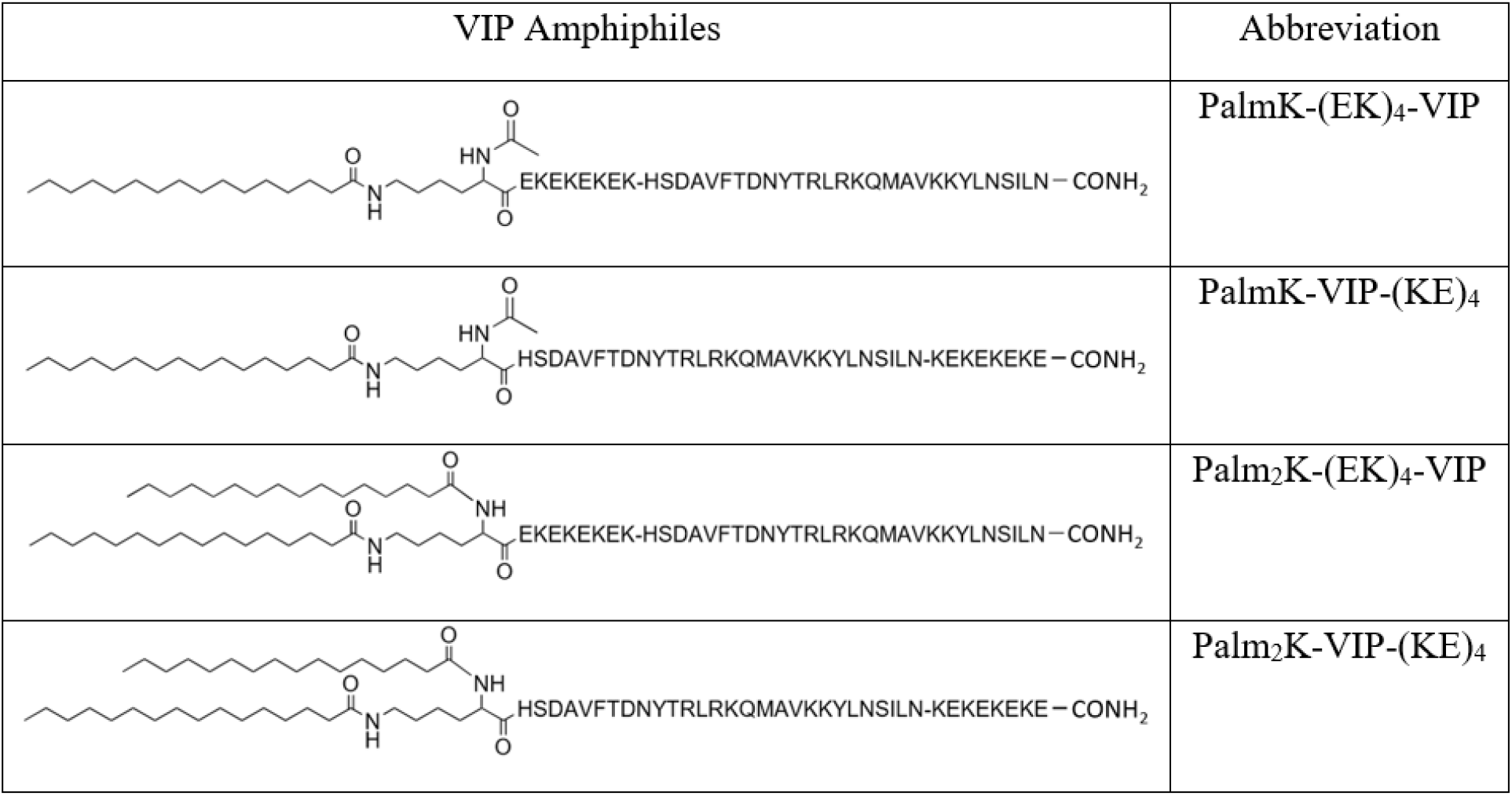
Chemical structure and abbreviated name of the triblock VIPAs studied.

After being synthesized on-resin, cleaved by trifluoroacetic acid, and purified by mass spectrometry controlled high-pressure liquid chromatography (LCMS), VIPAs were characterized by standard materials assessment techniques [25, 27, 28]. Critical micelle concentration (CMC) determination revealed the minimal solution concentration of PAs necessary to observe micelle formation (**Figure S1**). VIPA CMCs ranged from 0.0348 - 6.17 μM (**Table 2**) dependent on PA chemistry. Double lipid VIPAs (*i.e.*, Palm_2_K-(EK)_4_-VIP and Palm_2_K-VIP-(KE)_4_) were found to have CMCs 1 - 2 orders of magnitude lower than that of single lipid VIPAs (*i.e.*, PalmK-(EK)_4_-VIP and PalmK-VIP-(KE)_4_). This is consistent with previous results that suggest the difference in the hydrophobicity between lipid(s) and peptide governs PA micellization [28, 29]. A more modest effect on CMC was observed with zwitterion-like peptide block location. When the zwitterion-like peptide block was further away from the lipid group (*i.e.*, external block - PalmK-VIP-(KE)_4_ and Palm_2_K-VIP-(KE)_4_), the CMC was 6 - 9 times lower than with the analogous VIPA where the zwitterion-like peptide block was next to the lipid group (*i.e.*, internal block - PalmK-(EK)_4_-VIP and Palm_2_K-(KE)_4_-VIP). This is likely caused by the zwitterion-like peptide block being much more hydrophilic than VIP. Specifically, the zwitterion-like peptide block will arrange itself on the exterior of the micelle whereas the aromatic and aliphatic amino acids in VIP will try to associate more with the lipid core. When the zwitterion-like peptide block is external, PA micellization is readily ordered along the length of the molecule due to the gradation of its hydrophobicity. In contrast, when the zwitterion-like peptide block is internal, the PA must bend to accommodate the desired conformation of the zwitterion-like peptide block being on the surface of the micelle (**Figure S2**). This kink causes the formation of flower-like micelles which are commonly found to have higher CMCs, consistent with our previous research [28]. Interestingly, the double lipid, internal zwitterion-like peptide block PA (*i.e.*, Palm_2_K-VIP-(KE)_4_) was found to have two CMCs with one at 0.0348 μM and another at 0.337 μM. This result suggests the micelles may be going through a structural transition [30], for which further study by transmission electron microscopy (TEM) was warranted.

**Table 2.** Critical micelle concentration (CMC) of VIPAs.

| VIP amphiphile type | CMC (μM) | 2<sup>nd</sup> CMC (μM) |
| --- | --- | --- |
| PalmK-(EK)<sub>4</sub>-VIP | 6.17 | — |
| PalmK-VIP-(KE)<sub>4</sub> | 0.970 | — |
| Palm<sub>2</sub>K-(EK)<sub>4</sub>-VIP | 0.440 | — |
| Palm<sub>2</sub>K-VIP-(KE)<sub>4</sub> | 0.0348 | 0.337<sup>a</sup> |

a) The 2<sup>nd</sup> CMC was observed in three experimental repeats.

TEM was conducted on nano-tungsten negatively-stained VIPAs deposited on carbon-coated copper grids via rapid solvent wicking. Single-lipid VIPAs (*i.e.*, PalmK-(EK)_4_-VIP and PalmK-VIP-(KE)_4_) formed into long cylindrical micelles ranging from 0.2 - 7 μm in length, whereas double-lipid VIPAs (*i.e.*, Palm_2_K-(EK)_4_-VIP and Palm_2_K-VIP-(KE)_4_) generated short cylinders 30 - 400 nm in length (**Figure 1**). Furthermore, the longer single-lipid VIPAM cylinders were able to also associate with one another leading to elongated braid-like micellar aggregates (**Figure S3**). In contrast, the shorter double-lipid VIPAM cylinders did not appear to participate in micellar aggregation. These results illustrate that hydrophobicity was primarily responsible for VIPAM size and structure while zwitterion-like peptide block had a negligible impact on these properties. Interestingly, Palm_2_K-VIP-(KE)_4_ was found to have formed only spherical micelles approximately 5 - 20 nm in diameter at 0.1 μM, but predominantly cylindrical micelles at 1 μM (**Figure S4**). This suggests a sphere-to-cylinder transition event [30, 31] occurring for these PAs between these concentrations which is supported by a second CMC observed at 0.337 μM for this PA chemistry (**Table 2** and **Figure S1**). Interestingly, no such sphere-to-cylinder transition was observed for any of the other PA chemistries. Surprisingly, VIP at the highest concentration (*i.e.*, 100 μM) was found to aggregate into spherical nanoparticles (**Figure S5**), whereas no particle formation was observed at lower concentrations (*data not shown*). While VIPAM structure will play a key role in their interactions with cells, their surface charge will almost definitely be important in these interactions as well. Therefore, zeta potential of VIPAMs was measured to characterize the impact VIPA chemical structure has on micellar surface charge.

**Figure 1.**
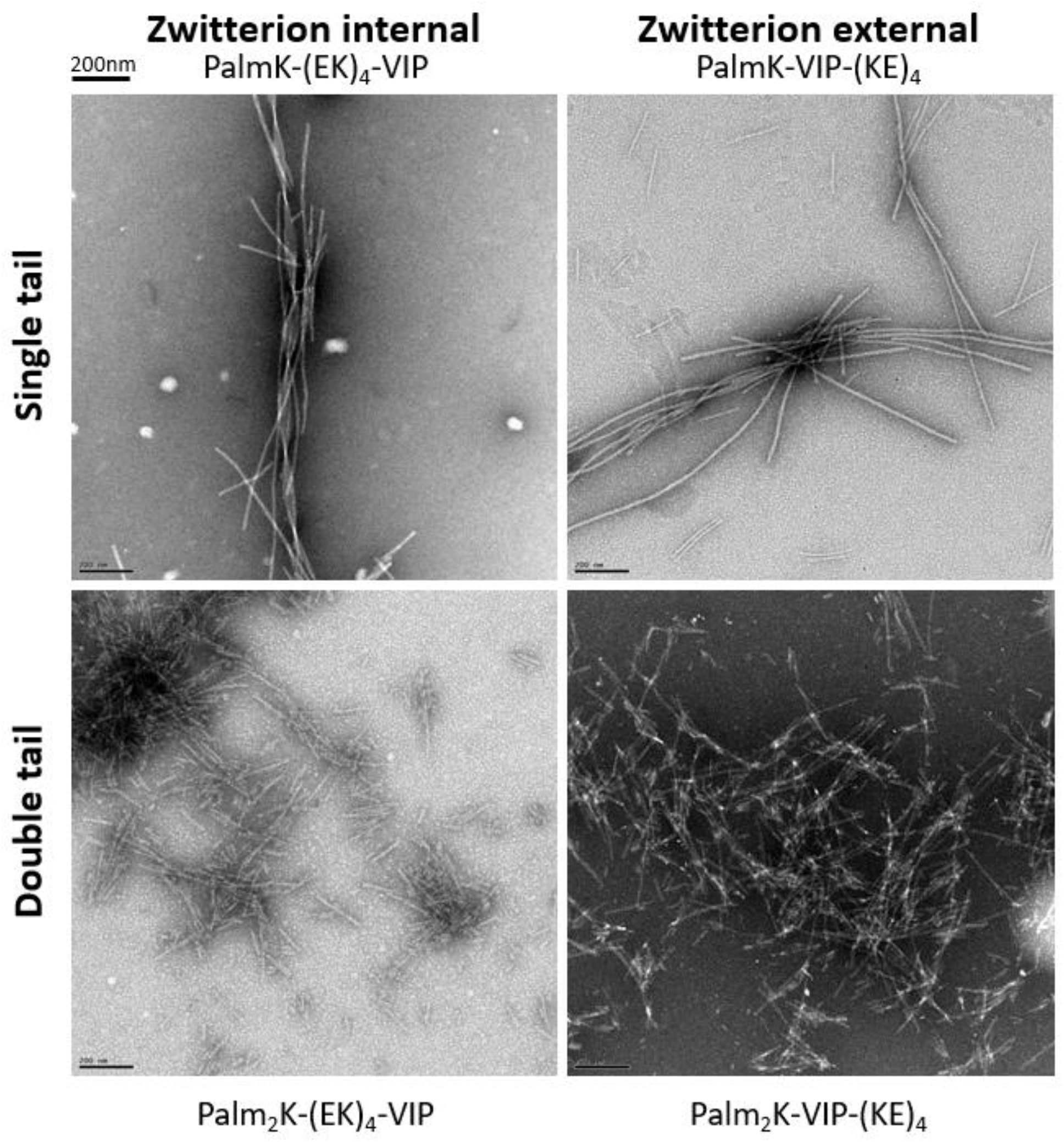
Morphology of VIP amphiphile micelles at 100 μM in phosphate buffered saline (PBS) as captured by transmission electron microscopy (TEM). All micrographs were taken at a magnification of 12,000 x with the scale bar for all images representing 200 nm.

VIPAM surface charge (*i.e.*, Zeta potential) was characterized by a DLS Zetasizer in deionized, distilled water (ddH_2_O) at a concentration of 100 μM (**Table 3**). Zwitterion-like peptide block position was found to exert the strongest influence on VIPAM surface charge. Internal zwitterion-like peptide block VIPAMs possessed higher positive surface charge than just VIP alone, whereas external zwitterion-like peptide block VIPAMs were less cationic than VIP by itself. This data suggests the organization of the VIPA peptide blocks within the micelles is heavily influenced by their location within the amphiphile. Specifically, the flower-like micellization of internal zwitterion-like peptide block VIPAMs folds the positively charged VIP between the zwitterion-like peptide block and the lipid core. To charge balance this region, the zwitterion-like peptide block can orient its anionic glutamic acid side chains to facilitate intra-amphiphile or intra-micellar charge complexation. This would result in the cationic lysine side chains of the zwitterion-like peptide block being presented on the micellar surface yielding the higher Zeta potential observed (**Figure S2**). In contrast, external zwitterion-like peptide block VIPAMs would not undergo this amphiphile folding constraint limiting the need for internal charge complexation yielding a surface more balanced between the glutamic acid and lysine side chains of the zwitterion-like block resulting in the more neutral Zeta potential observed. Though all four VIPAs were able to self-assemble into micelles capable of preventing VIP molecules from diffusing away from one another thereby increasing the potential for local payload concentration, they possessed varying surface charges which may impact their interactions with cells. Therefore, the combined effect of micelle structure and surface charge on modulating the bioactivity of VIP was studied using antigen presenting cells.

**Table 3.** Zeta potential of VIP and VIPAs.

| VIP Amphiphile Micelle Type | Zeta potential (mV) |
| --- | --- |
| VIP | 6.82 ± 0.67 |
| PalmK-(EK)<sub>4</sub>-VIP | 9.27 ± 0.084 |
| PalmK-VIP-(KE)<sub>4</sub> | 2.26 ± 0.68 |
| Palm<sub>2</sub>K-(EK)<sub>4</sub>-VIP | 10.6 ± 0.47 |
| Palm<sub>2</sub>K-VIP-(KE)<sub>4</sub> | 3.79 ± 0.085 |

### VIPA and VIPAM impact on Mφ health and inflammation modulation

Murine Mφs (100,000 cells) were activated with 0.1 μg/mL lipopolysaccharide (LPS) and treated with various VIPA concentrations (*i.e.*, 0.1 μM, 1 μM, 10 μM, or 100 μM). Mφs given only the PBS vehicle (*i.e.*, no LPS) served as a negative control and cells exposed to LPS alone acted as a positive control for cell count and tumor necrosis factor α (TNF-α) secretion analysis. After 6 hours and 24 hours of incubation, Mφs were lysed using a 1% Triton X-100 solution and characterized by the Quant-IT PicoGreen dsDNA assay to investigate VIPA cytotoxicity (**Figure 2**). Before lysis, media supernatants were collected and VIPA anti-inflammatory effects were characterized by TNF-α secretion as determined by an enzyme-linked immunosorbent assay (ELISA) (**Figure 3**).

**Figure 2.**
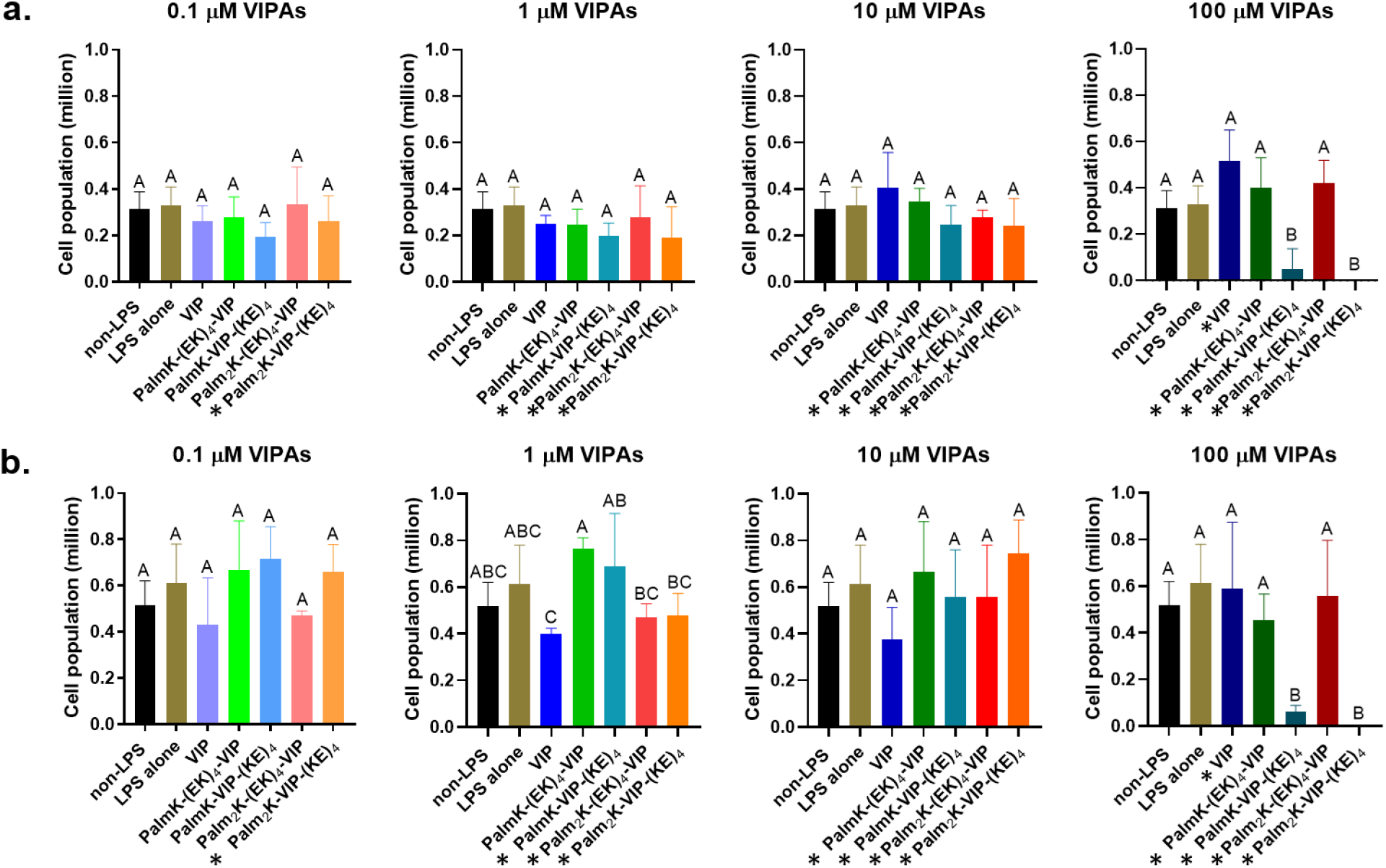
Effect of VIPAs on LPS-activated Mφ cell health after 6 hours (a) and 24 hours (b) of treatment (starting concentration - 100,000 cells per group). In each graph, groups possessing the same letter have no statistically significant difference (p > 0.05). The “*” sign in front of the name of a VIPA formulation indicates it is above the CMC, therefore micelles were present.

**Figure 3.**
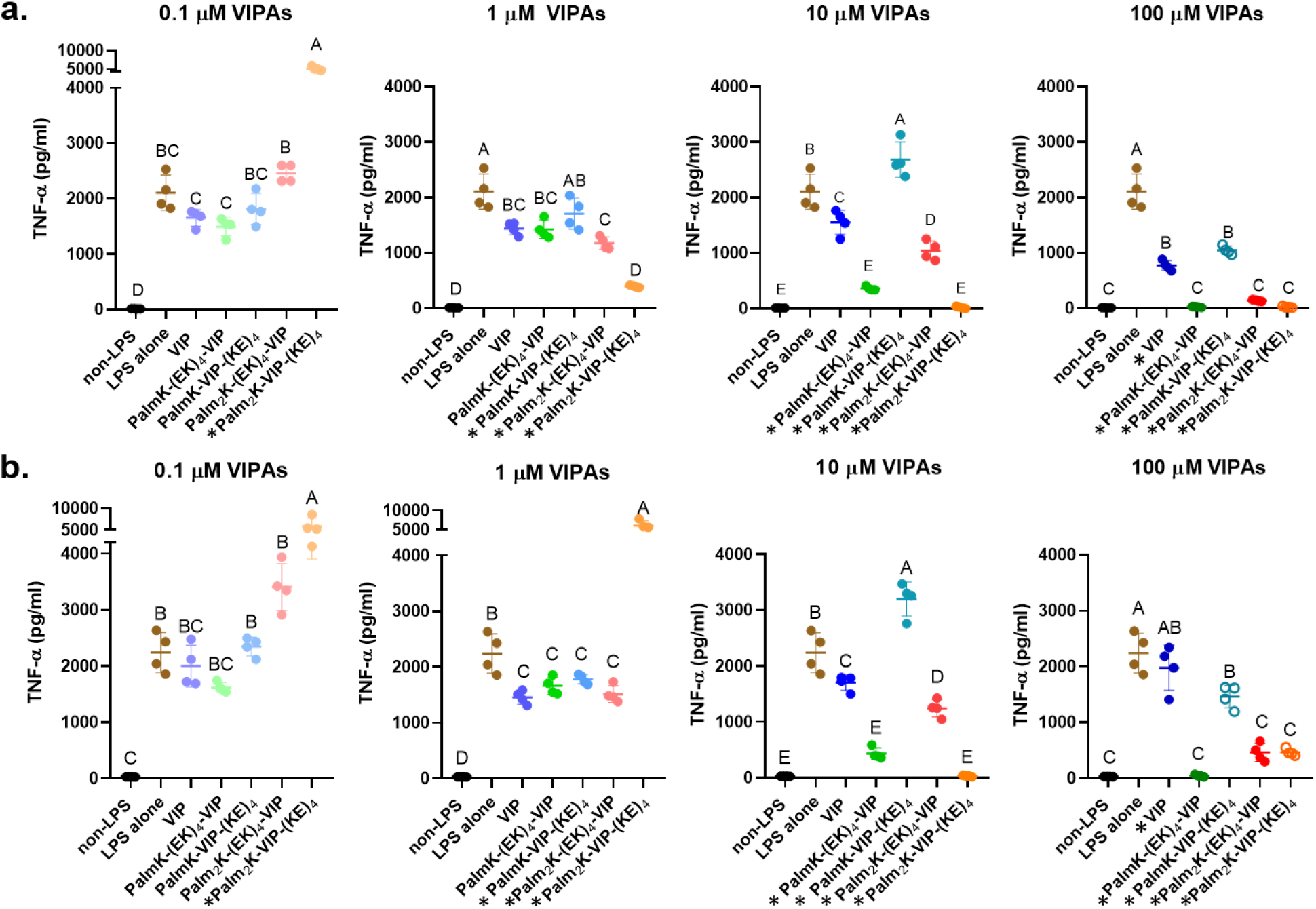
Effect of VIPAs on LPS-activated Mφ TNF-α secretion after 6 hours (a) and 24 hours (b) of treatment. In each graph, groups possessing the same letter have no statistically significant difference (p > 0.05). The “*” sign in front of the name of a VIPA formulation indicates it is above the CMC, therefore micelles were present. The open dot data points (i.e., 100 μM PalmK-VIP-(KE)_4_ and 100 μM Palm_2_K-VIP-(KE)_4_) correspond to cytokines secreted from cells in poor health due to exposure to high VIPA concentration.

All VIPA formulations at 0.01 - 10 μM showed no immediate cytotoxicity in 6 hours (**Figure 2a**), but the 100 μM doses of both external zwitterion-like peptide block VIPAMs (*i.e.*, PalmK-VIP-(KE)_4_ and Palm_2_K-VIP-(KE)_4_) induced significant cell death by 24 hours (**Figure 2b**). As the internal zwitterion-like peptide block VIPAMs (*i.e.*, PalmK-(KE)_4_-VIP and Palm_2_K-(KE)_4_-VIP) do not possess similar high concentration toxicity, the data suggests that VIPA chemical structure influences its biocompatibility at high concentrations. Specifically, it is conceivable that cell cytotoxicity at 100 μM for some of the formulations was caused by osmotic stress since previous studies that employed particles with moderately positive surface charge showed them to be more effectively phagocytosed by macrophages than their more neutral counterparts [32, 33]. This suggests the more cationic internal zwitterion-like peptide block VIPAMs would be more easily internalized by macrophages when compared with the more neutral external zwitterion-like peptide block VIPAMs. A greater intracellular VIPAM concentration would mitigate the osmotic pressure difference across the cell membrane allowing it to maintain its integrity. In contrast, the imbalance created by the less cationic external zwitterion-like peptide block VIPAMs would lead to a large flux of water from the inside of the cells out leading to the observed cell cytotoxicity. No further cytotoxicity for other VIPA groups was observed in the first 24 hours of incubation indicating a quickly developing imbalance in osmotic pressure may be a primary reason for the observed behavior.

TNF-α secretion from VIPA-treated LPS-induced Mφs was then analyzed to determine the anti-inflammatory behavior of the various formulations. Interestingly, the anti-inflammatory effects of VIPAs were found to be time, concentration, and formulation dependent (**Figure 3**). At 6 hours, VIP peptide alone induced mild to moderate TNF-α suppression in LPS-activated Mφs in a concentration-dependent fashion (**Figure 3a**). VIPAs, when present as individual amphiphiles at this time point, showed little to no ability to modulate the TNF-α suppression capacity of the incorporated VIP peptide. In contrast, VIPA micellization was found to significantly alter VIP anti-inflammatory effects in macrophages. Micelles comprised of PalmK-(EK)_4_-VIP (10 - 100 μM) and Palm_2_K-(EK)_4_-VIP (1 - 100 μM) induced modest to significant TNF-α suppression when compared to VIP peptide alone. In contrast, PalmK-VIP-(KE)_4_ micelles at high concentration (i.e., 10 μM) were found to significantly increase macrophage TNF-α secretion, an effect not observed at lower concentration (*i.e.*, 1 μM). Interestingly, Palm_2_K-VIP-(EK)_4_ micelles were found to exert a unique concentration-dependent influence. At low concentration (0.1 μM), they facilitated considerable enhancement in TNF-α expression, whereas at higher concentrations (1 - 10 μM), they caused significant suppression of TNF-α secretion. The anti-inflammatory effects of external zwitterion-like peptide block micelles (*i.e.*, PalmK-VIP-(KE)_4_ and Palm_2_K-VIP-(KE)_4_) at high concentration (100 μM) were obfuscated by the low number of cells present. Therefore, TNF-α secretion was also normalized by cell count (**Figure S6**), which clearly demonstrated the pro-inflammatory effect of PalmK-VIP-(KE)_4_ at this concentration. The immunomodulatory properties observed for the micelles appeared to all correlated to their size and shape.

TNF-α suppressive VIPAM formulations (*i.e.,* 10 - 100 μM PalmK-(EK)_4_-VIP, 1 - 100 μM Palm_2_K-(EK)_4_-VIP, and 1 - 100 μM Palm_2_K-VIP-(KE)_4_) were all found to be cylinders that were 30 nm - 3 μm in size. In contrast, TNF-α enhancing VIPAM formulations were either small spherical micelles (0.1 μM Palm_2_K-VIP-(EK)_4_) or elongated braided micelles (10 - 100 μM PalmK-VIP-(KE)_4_). While at opposite ends of the particle size scale, the pro-inflammatory effects of products with similar dimensions as these formulations has been previously published. In specific, macrophages have been shown to bind small, spherical nanoparticles much faster than cylindrical nanoparticles [34], which likely plays an important role in their enhanced phagocytosis [35]. Additionally, particle-triggered inflammation is a size-dependent process where the larger the particle size, the higher potential to induce inflammation [36]. Fiber-like microparticles with longer lengths (> 7 μm) have been found to induce high levels of TNF-α expression [37]. As the cell receptor for VIP (*i.e.*, VPAC) is transmembrane in nature and presented externally on the cell surface, cylindrical micelles large enough to slow their internalization, but small enough to avoid over-activating TNF-α expression makes them ideal for enhancing VIP-mediated anti-inflammatory effects. Interestingly, TNF-α secretion barely changed in the LPS treated group from 6 hours to 24 hours implying LPS-induced inflammation in Mφs is a rapid process (**Figure 3b**). During this longer incubation period, all relative changes in TNF-α secretion were caused solely by prolonged exposure to various VIP formulations. Interestingly, 10 μM and 100 μM VIPAMs, except for PalmK-VIP-(KE)_4_, induced similar TNF-α suppression effects at 24 hours as was observed with the same formulations and concentrations at 6 hours. Specifically, 24 hours of 1 μM Palm_2_K-VIP-(KE)_4_ incubation induced Mφs to produce much more TNF-α in contrast to the suppressive behavior observed with the same concentration and formulation at 6 hours. This behavior may be attributable to Mφs processing enough VIPAMs from solution that their concentration dropped below the higher CMC triggering cylindrical to spherical micelle transition inducing similar behavior to what was observed for 0.1 μM Palm_2_K-VIP-(KE)_4_ at both 6 hours and 24 hours. The influence of nanoparticle shape may also explain the change in TNF-α inducing behavior of VIP at 100 μM from 6 to 24 hours, where cytokine secretion was suppressed at the early time point, but increased to the LPS-stimulus alone level by the later time point. This result may suggest there is a competing effect between the pro-inflammatory effect of spherical particles like what was observed with low concentrations for Palm_2_K-VIP-(KE)_4_ micelles and the anti-inflammatory effect the high dose of VIP exerts. These results provide considerable promise for the potential internal zwitterion-like peptide block VIPAMs (*i.e.*, PalmK-(EK)_4_-VIP and Palm_2_K-(EK)_4_-VIP) have as prolonged anti-inflammatory therapeutics. That being said, their impact on other APCs like DCs is also of importance in determining their translatability.

### VIPA and VIPAM impact on DC health and inflammation modulation

Murine bone-marrow derived dendritic cells (DCs) were activated with 0.1 μg/mL LPS and treated with various VIPA concentrations (*i.e.*, 0.1 μM, 1 μM, 10 μM, or 100 μM). DCs exposed only to the PBS vehicle (*i.e.*, no LPS) or LPS alone acted as negative and positive controls, respectively. All groups were assessed for cell viability, cytokine secretion, and cell surface marker expression after 6-hour and 24-hour incubation periods. Cytokine analysis was performed using the same method employed for Mφs though the cytokine CCL22 was analyzed by ELISA in addition to TNF-α. After supernatant collection, DCs were detached using enzyme-free dissociation buffer and stained by propidium iodide for cell viability characterization and by fluorescent antibodies specific for MHC II and CD86 before assessment by flow cytometry to determine the impact VIPAs have on DC activation.

All VIPA formulations at 0.1 - 10 μM and almost all VIPA formulations at 100 μM were found to be non-toxic to DCs after 6 hours of incubation (**Figure 4a**). Palm_2_K-VIP-(KE)_4_ at 100 μM induced a slight decrease in cell viability causing over 30% of cells to die. PalmK-VIP-(KE)_4_ at 100 μM was found to be quite toxic with less than a third of DCs being viable post-exposure. At 24 hours, both external zwitterion-like block VIPAs (*i.e.,* PalmK-VIP-(KE)_4_ and Palm_2_K-VIP-(KE)_4_) at 100 μM and PalmK-VIP-(KE)_4_ at 10 μM induced death in more than half of the DCs they were incubated with (**Figure 4b**). These results are similar to what was observed with Mφs providing evidence that this behavior is APC cell-type independent and potentially consistent with the previously mentioned osmotic stress theory.

**Figure 4.**
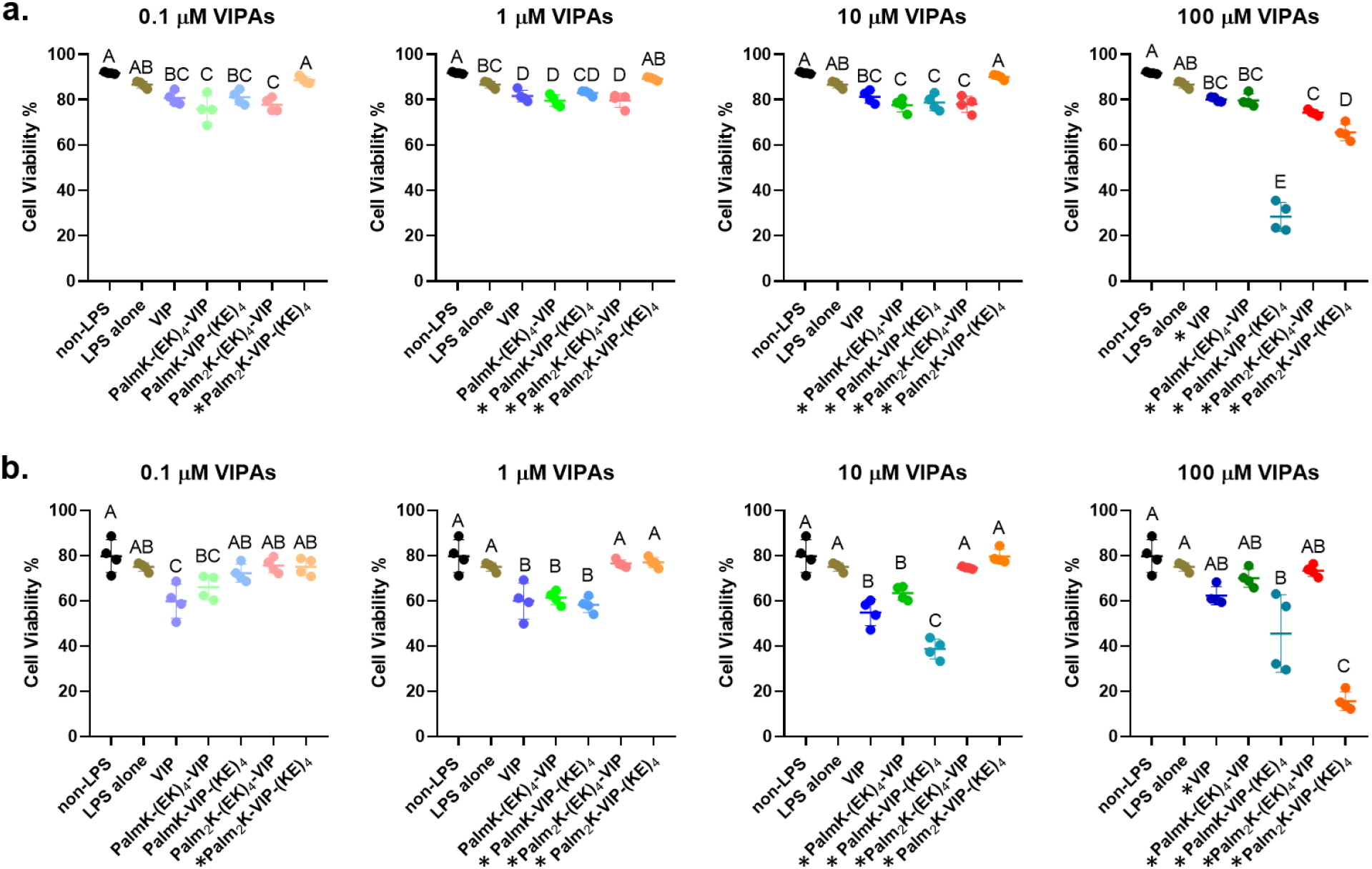
Effect of VIPAs on LPS-activated DC cell health after 6 hours (a) and 24 hours (b) of treatment. In each graph, groups possessing the same letter have no statistically significant difference (p > 0.05). The “*” sign in front of the name of a VIPA formulation indicates it is above the CMC, therefore micelles are present.

Enhancement of TNF-α suppression by VIPAs in DCs was found to be time, concentration, and formulation dependent like what was observed for Mφs, though somewhat different aspects of these factors influenced the anti-inflammatory effects of the formulations (**Figure 5**). Concentration-dependent TNF-α inhibition of VIP was observed at 6 hours (**Figure 5a**). VIP alone was able to decrease TNF-α secretion from LPS-stimulated DCs slightly at 10 μM and more extensively at 100 μM at this time point, whereas all VIPA formulations potentiated mild to significant TNF-α suppression greater than VIP alone in a concentration-dependent fashion from 1 - 100 μM. While some overall similarities were observed between Mφ and DC inflammation mediation by VIPAs, micelle shape appeared to be less influential for DCs, for which just having VIPA-based micelles present was found to be the most important factor at 6 hours. In contrast, formulation-determined shape and size effects started to influence TNF-α secretion in DCs between 6 and 24 hours (**Figure 5b**). First, a major difference between LPS-stimulation of Mφs and DCs was found during this time period. Whereby Mφs maintained high levels of extracellular TNF-α at both 6 and 24 hours due to exposure to the pro-inflammatory LPS stimulus, DCs saw a significant decrease from high levels at 6 hours to much lower levels at 24 hours [38]. Compared to the lower baseline level of LPS-induced TNF-α secretion at 24 hours, prolonged exposure of DCs to small spherical micelles / nanoparticles (*i.e.,* 0.1 μM Palm_2_K-VIP-(KE)_4_ and 100 μM VIP) and elongated cylindrical micelles (*i.e.,* 1 μM and 10 μM PalmK-VIP-(KE)_4_) yielded slightly to modestly increased cytokine production. In contrast, all cylindrical micelle formulations 30 nm – 3 μm in length continued to facilitate TNF-α secretion suppresion through 24 hours. This discovery illustrated that rather than being completely unresponsive to micelle shape and size, DCs have a more delayed response to pro-inflammatory micelles when compared to Mφs. This phenomenon may be related to the different environmental sampling mechanisms Mφs and DCs employ when activated. Specifically, DCs downregulate their capacity for micropinocytosis and switch to more receptor-mediated endocytosis during maturation in contrast to activated Mφs which still favor macropinocytosis [39, 40]. As receptor-mediated endocytosis is a slower internalization mechanism, VIPA-based delivery of VIP to the DC surface allows for stimulation of the cognate cell-surface receptor (*i.e.*, VPAC for VIP) in a size and shape independent manner and is the dominant immunomodulatory factor at 6 hours. By 24 hours, internalization can have proceeded to the point where size and shape dependent factors also influence DC immunomodulation. Interestingly, the total extracellular quantity of TNF-α present around LPS-activated DCs decreased by nearly 90% from 6 to 24 hours indicating the cells must have internalized and processed the secreted TNF-α. This process is known to promote other events in DCs including maturation [41], apoptosis [42], and additional cytokine production including IL-6, IL-8, and CCL22 [43–45]. With this in mind, CCL22 production was analyzed by ELISA (**Figure S7**) due to its ability to recruit regulatory T cells [46, 47] and since it has been reported to be inducible by both TNF-α [45] and VIP [48]. Interestingly, we found that the CCL22 production was induced by LPS stimulation, and its production was promoted between 6 hours to 24 hours. This enhancement of CCL22 secretion aligns with the drop in secreted TNF-α present between the time points. Moreover, VIP or VIPA groups that induced pro-inflammatory effects as measured by TNF-α secretion (*i.e.,* 0.1 μM Palm_2_K-VIP-(KE)_4_, 1μM and 10 μM PalmK-VIP-(KE)_4_, and 100 μM VIP), showed enhanced CCL22 expression. In congruence with this relationship, groups which suppressed TNF-α generation (*i.e.*, 0.1 μM – 100 μM PalmK-(EK)_4_-VIP and Palm_2_K-(EK)_4_-VIP, 1 μM −100 μM Palm_2_K-VIP-(KE)_4_, and 1 μM and 10 μM VIP) also reduced CCL22 production. While some published results have shown VIP is able to increase CCL22 expression from LPS-activated DCs [46, 48], these studies focused on longer incubation times of at least 48 hours. In fact, no previously published work to date has explored in depth the relationship between VIP and CCL22 upregulation within the first 24 hours. Our results suggest that CCL22 secretion from DCs early in the maturation process is more regulated by the initial presence of a high extracellular TNF-α concentration than VIP or VIPA concentration. In addition to altering DC cytokine production, VIP has been found to modulate adaptive immunity activation by down-regulating DC surface markers (*i.e.*, MHC II and CD86) that play a role in T cell stimulation [26, 49]. Therefore, these proteins were characterized to explore the potential VIPAs have to impact this important step in adaptive immune response induction.

**Figure 5.**
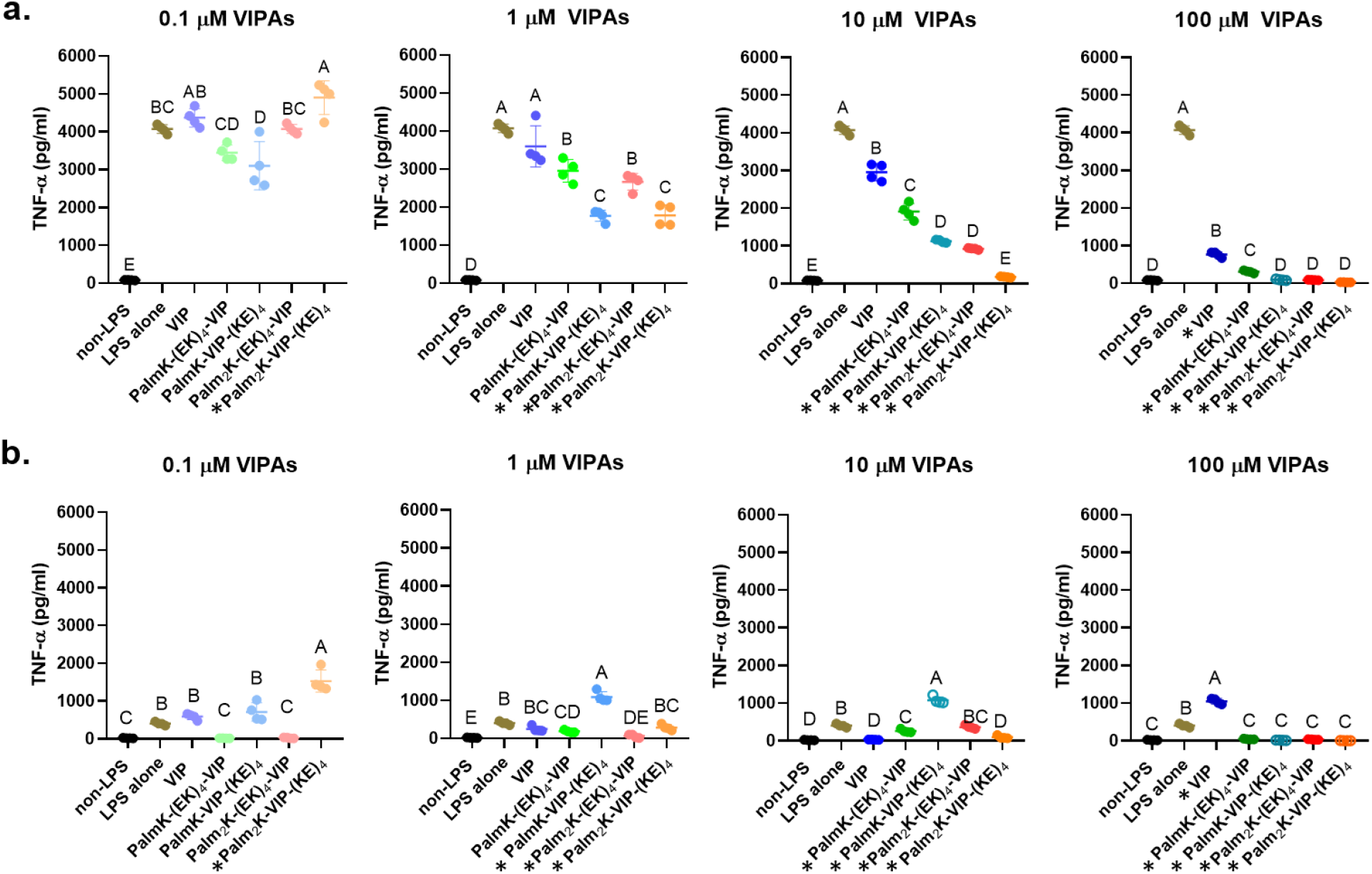
Effect of VIPAs on LPS-activated DC TNF-α secretion after 6 hours (a) and 24 hours (b) of treatment. In each graph, groups possessing the same letter have no statistically significant difference (p > 0.05). The “*” sign in front of the name of a VIPA formulation indicates it is above the CMC, therefore micelles were present. Data dots with no fill indicates this group has more than 50% non-viable cell.

### VIPA and VIPAM impact on DC surface marker expression

MHC II is a protein complex located on the surface of APCs which binds peptide fragments derived from antigens for presentation to CD4^+^ T cells [50]. CD86 is a co-stimulatory receptor modulated by APC activation that works synergistically with MHC II to guide T cell recognition and activation [51]. As both MHC II and CD86 are necessary for adaptive immunity activation, the percentage of MHC II^+^CD86^+^ DCs was determined (**Figure 6**). DC cell surface marker expression was found to be highly dependent on incubation time and VIPA formulations. In general, VIP and most VIPAs were found to have limited influence on the MHC II^+^CD86^+^ DC population at 6 hours, however, more distinct impacts on MHC II^+^CD86^+^ DCs regulation were observed at 24 hours. Interestingly, MHC II and CD86 expression on DCs at 24 hours greatly aligned with TNF-α and CCL22 secretion at 24 hours supporting the important relationship between cytokine production and cell surface marker expression [52]. Specifically, most of the groups that suppressed TNF-α production (*i.e.,* 1 - 10 μM VIP, 0.1 - 100 μM PalmK-(EK)_4_-VIP, 0.1 μM - 1 μM Palm_2_K-(EK)_4_-VIP, and 10 μM Palm_2_K-VIP-(KE)_4_) also downregulated MHC II and CD86 surface markers. In contrast, most groups that were unable to suppress TNF-α generation (*i.e.,* 100 μM VIP, 0.1 – 1 μM PalmK-VIP-(KE)_4_, and 0.1 μM Palm_2_K-VIP-(KE)_4_) failed to downregulate MHC II and CD86. Among all VIPA formulations, internal zwitterion-like peptide block VIPAs (*i.e.,* PalmK-(EK)_4_-VIP and Palm_2_K-(EK)_4_-VIP) had the most interesting impact on MHC II and CD86 expression on DCs. PalmK-(EK)_4_-VIP induced considerable cell surface marker suppression at all concentrations (*i.e*., 0.1 - 100 μM) at 24 hours, specifically decreasing the population of LPS-stimulated MHC II^+^CD86^+^ DCs by 40% to 80% depending on concentration. Palm_2_K-(EK)_4_-VIP was also very potent in suppressing cell surface marker presentation, but interestingly only at lower concentrations (*i.e.,* 0.1 and 1 μM) inducing a decrease in MHC II^+^CD86^+^ DCs by approximately 55% and 70% at 6 and 24 hours, respectively, when compared to LPS-stimulated DCs. In contrast, high concentrations (*i.e.*, 10 and 100 μM) of Palm_2_K-(EK)_4_-VIP showed negligible impact at 6 hours and a 30% - 40% increase at 24 hours in the quantity of MHC II^+^CD86^+^ DCs over the LPS-stimulated control cells. These unique results may be related to competition between various influences VIPAMs exert on DCs. Specifically, the more rapid onset of effects by Palm_2_K-(EK)_4_-VIP as compared to PalmK-(EK)_4_-VIP may be due to their more hydrophobic double lipid region being able to insert into the cell bilayer more readily [53] helping to better facilitate quicker engagement of VPAC. While helpful for facilitating on-target VIP effects, this may be a double-edged sword since it is likely to also lead to greater quantities of nanoparticles associating with the cells possibly causing further DC activation in a concentration-dependent fashion similarly to what has been observed with other nanoparticles [33, 54, 55]. At the higher concentrations of Palm_2_K-(EK)_4_-VIP, these competing cell surface marker suppression and enhancement effects may be balanced at the earlier time point (*i.e.*, 6 hours), but the nanoparticle-mediated activation impact is greater by the later time point (*i.e.*, 24 hours). At lower concentrations of Palm_2_K-(EK)_4_-VIP, nanoparticle accumulation may not occur in great enough quantity to trigger this effect, so VIP-mediated suppression is able to dominate.

**Figure 6.**
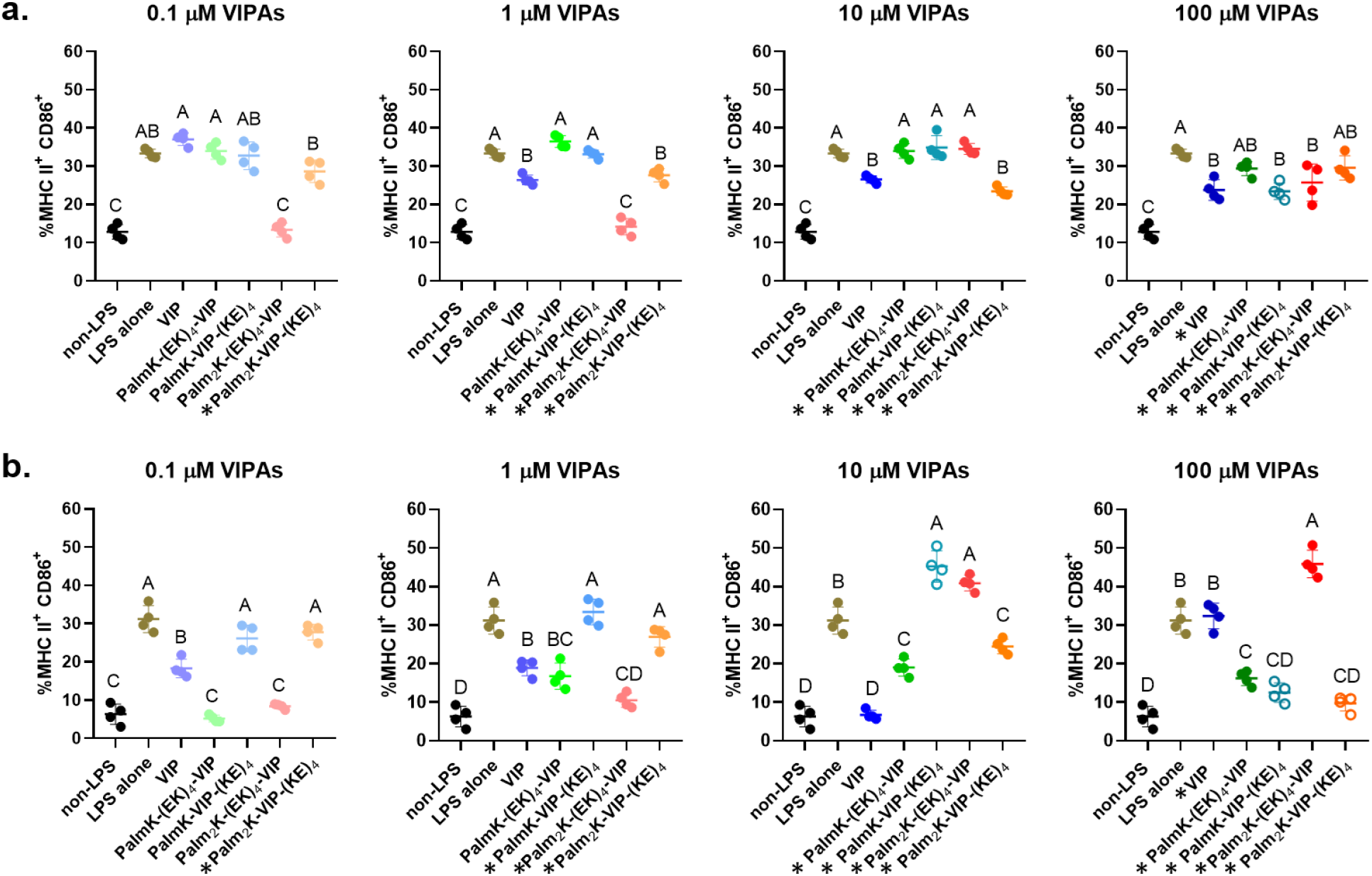
VIPAs treated MHC II^+^CD86^+^ dendritic cells population percentage after 6 hours (a) and 24 hours (b) treatment. Data was collected by flow cytometry and analyzed by FlowJo software. In each graph, groups possessing the same letter have no statistically significant difference (p > 0.05). The “*” sign in front of the name of the VIPA formulation indicates it is above the CMC and micelles have been formed. Data dots with no fill indicates groups with less than 50% viable cells.

## Conclusion

The studies conducted provide valuable insight into the impact micelle hydrophobic domain size and zwitterion-like peptide block location have on nanoparticle physical properties as well as their ability to potentiate or inhibit the function of an incorporated immunomodulatory peptide (*i.e.*, VIP). The concentration at which VIPAs formed micelles (*i.e.*, CMC) was influenced by both of these properties. In contrast, the hydrophobic domain governed VIPA micelle size and zwitterion-like peptide block location-controlled nanoparticle surface charge. VIPAs and VIPAMs were biocompatible regardless of formulation at most concentrations with more positively charged VIPAMs being less toxic to APCs at very high concentrations than more neutral formulations. Overall, the anti-inflammatory effects of VIPAs and VIPAMs were found to be time, concentration, and formulation dependent, but relatively independent of APC type. Cylindrical VIPAMs 30 nm - 3 μm in length were shown to possess the best anti-inflammatory behavior, whereas spherical micelles or longer cylindrical micelles and aggregates actually induced pro-inflammatory responses. Moreover, this initial screening yielded a promising lipidated VIP formulation (*i.e.,* PalmK-(EK)_4_-VIP) with more robust anti-inflammatory effects than unmodified VIP (**Figure 7**). Further formulation-dependent nuances were observed for VIPAMs within this size range as the single lipid, internal zwitterion-like peptide block amphiphile (PalmK-(EK)_4_-VIP) and the double lipid, internal zwitterion-like peptide block amphiphile (Palm_2_K-(EK)_4_-VIP) were found to have interesting time-dependent and concentration-dependent immunomodulatory effects on DC activation. This work details the importance many factors including chemical structure, hydrophobicity, micelle size, and surface charge have on peptide amphiphile micelle structure and bioactivity. While encouraging, further research is still needed to better understand the relationship micelle association and uptake by APCs have on their ability to enhance the function of incorporated VIP as well as influence adaptive immunity activation.

**Figure 7.**
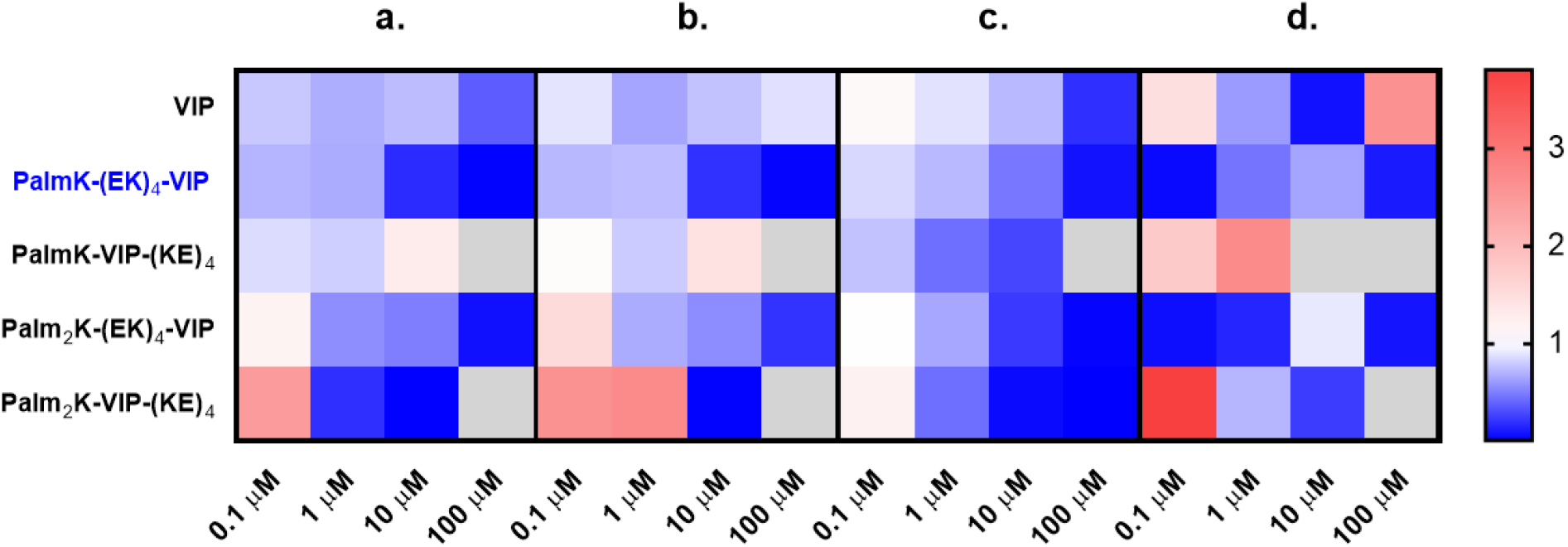
A heatmap representing fold changes of TNF-α expression in Mφs at 6 hours (a) and 24 hours (b), and in DCs at 6 hours (c) and 24 hours (d) relate LPS alone control groups to those treated with LPS and VIP or VIP formulations at different concentrations. Gray cells indicate treatment conditions where poor cell health was observed.

## ASSOCIATED CONTENT

### Supporting Information

Supporting Information is available free of charge at ***Link***. Materials and methods, critical micelle concentration graphs, details on the structure of VIPA micelles, and additional data on cytokine secretion of Mφ and DCs can all be found within this document.

### Authors

**Xiaofei Wang** – Department of Biomedical, Biological, and Chemical Engineering, University of Missouri, Columbia, MO 65211

**Rui Zhang** – Department of Biomedical, Biological, and Chemical Engineering, University of Missouri, Columbia, MO 65211

**Bryce D. Lindaman** – Department of Biomedical, Biological, and Chemical Engineering, University of Missouri, Columbia, MO 65211

**Catlin N. Leeper** – Department of Biomedical, Biological, and Chemical Engineering, University of Missouri, Columbia, MO 65211

**Adam G. Schrum** – Departments of Molecular Microbiology & Immunology, Surgery, and Biomedical, Biological & Chemical Engineering, University of Missouri, Columbia, MO 65211

## Acknowledgment

We thank Dr. Fabio Gallazzi of the Molecular Interactions Core at the University of Missouri for purifying the VIP and VIPAs used in this research. This work was funded by the Office of the Assistant Secretary of Defense for Health Affairs and the Defense Health Agency J9, Research and Development Directorate through the Reconstructive Transplant Research Program under Award No. W81XWH-17-1-0596. Opinions, interpretations, conclusions, and recommendations are those of the author and are not necessarily endorsed by the Department of Defense.

## Notes

### Competing Interest Statement

The authors have declared no competing interest.

