## Supplementary Information for "Vasoactive Intestinal Peptide Amphiphile Micelle Chemical Structure and Hydrophobic Domain Influence Immunomodulatory Potentiation"

### Materials and Method

**Solid Phase Peptide Synthesis (SPPS) and Lipidation.** Vasoactive intestinal peptide (VIP - Ac-HSDAVFTDNYTRLRKQMAVKKYLNSILN-$CONH_2$), Fmoc-EKEKEKEK-VIP-CONH-Resin, and Fmoc-VIP-KEKEKEKE-CONH-Resin were purchased from Synpeptide Co., Ltd, China. Fmoc-Lys(Fmoc)-OH and Fmoc-Lys(ivDde)-OH were acquired from Novabiochem and 2-(1H-benzotriazol-1-yl)-1,1,3,3-tetramethyluronium hexafluorophosphate (HBTU) and 1-hydroxybenzotriazole hydrate (HOBt hydrate) were purchased from AK Scientific, Inc. and AnaSpec, Inc., respectively. Piperidine, dimethylfuran (DMF), 1-methyl-2-pyrrolidinone (NMP), N,N-diisopropylethylamine (DIEA), acetic anhydride, hydrazine monohydrate, thioanisole (TA), trifluoroacetic acid (TFA), triisopropylsilane (TIS), phenol, ethanedithiol (EDT), and diethyl ether were bought from Millipore Sigma. A glass peptide reaction vessel was obtained from Chemglass Life Sciences.

Peptides were modified on amide resin employing solid phase synthesis in a glass vessel using Fmoc chemistry. The dry resin was rinsed in NMP for 2 hours with nitrogen gas bubbling. *Fmoc removal:* The Fmoc group was removed by 30-minute treatment with 25% piperdine in DMF carried out twice. *Amino acid conjugation:* Fmoc-Lys(ivDde)-OH (1 eq.) or Fmoc-Lys (Fmoc)-OH (1 eq.) was activated using uronium salts (HBTU, 4.2 eq., HOBt, 5 eq.) and DIEA (10 eq.) in NMP for 10 minutes and was conjugated on the N terminus of the peptide after 2 times of 1.5-hour coupling. *Lipidation:* Following Fmoc deprotection of either Fmoc-Lys(ivDde)-OH (for single lipid tail VIPA formulas) or Fmoc-Lys (Fmoc)-OH (for double lipid tails VIPA formulas) that had been added to the N terminus of the peptide, a palmitic acid or palmitic acids were added. *IvDde removal:* IvDde was removed by treating the single lipidated peptide formulations on resin with 2% hydrazine monohydrate in DMF for 20 minutes for a total of 6 times. The free amine was then acetylated through exposure to a solution of 5% acetic anhydride and 7% DIEA in DMF. *Peptide cleavage:* All peptides were cleaved from resin and their side chain protecting groups through a single-step reaction comprised of a 2-hour exposure to 87.5:2.5:2.5:2.5:2.5:2.5 TFA:TA:phenol:water:EDT:TIS. Cleaved peptide was precipitated in diethyl ether and washed with excess diethyl ether 5 times. All products synthesized were assessed by analytical high-pressure liquid chromatography (HPLC, Beckmann Coulter, Fullerton, CA) and purified by mass spectrometry-controlled liquid chromatography (LC-MS) using a C4 column (Milford, MA). The sequence of the four peptide amphiphiles are shown in Error! Reference source not found..

**Critical Micelle Concentration (CMC).** The hydrocarbon 1,6-diphenyl-1,3,5-hexatriene (DPH), a molecule that is weakly fluorescent in water, but strongly fluorescence when stacked in a hydrophobic domain, was utilized as an indicator for determining the critical micelle concentration (CMC). Serial diluting concentrations of peptide amphiphiles (316 μM to 0 μM) were made in 1 μM DPH in phosphate buffered saline (PBS) and allowed to incubate for at least 1 hour. Sample fluorescence was measured by a BioTek Cytation 5 fluorospectrophotometer (*ex*: 350 nm, *em*: 428 nm). The intersection point between trend lines for low fluorescence and rapidly increasing fluorescence was considered the CMC for the sample.

**Micelle Morphology Characterization.** The morphology of peptide amphiphile micelles were characterized by negative-stain aided transmission electron microscopy (TEM). A small amount (5 μL) of product solution (100 μM or other specified concentrations) was added to a carbon support TEM grid (200 mesh, Electron Microscopy Sciences). After 5 minutes of incubation, excessive solution was wicked away by filter paper and immediately followed with 5 μL of nanotungsten (Nanoprobes, Inc) solution added to the grid. After 3 minutes of incubation, grids were blotted dry and imaged with a JEOL JEM-1400 TEM at 120 kV. At least 3 different spots on each grid were analyzed for which the images presented are representative ones.

**Micelle Surface Charge Measurement.** Micelle surface charge was assessed by determining its zeta potential in zero charged medium. Product solutions (100 μM) were prepared in deionized distilled water and measured by a Malvern DLS Zetasizer.

**Cell Differentiation, Culture and Stimulation.** The initial anti-inflammatory experiments were conducted with mouse RAW 264.7 (macrophage-like cells - Mφs). Complete Dulbecco's Modified Eagle Medium (DMEM) was made by mixing DMEM (ThermoFisher), 10% fetal bovine serum (FBS, Sigma Aldrich), and 1% penicillin/streptomycin (ThermoFisher). Mφs were cultured in 100 mm diameter non-treated petri-dishes in complete DMEM at 37 °C and 5% $CO_2$ with media changes every two days until the cells became confluent. The cells were then seeded in non-treated 24-well plates at 0.1 million cells per well. After overnight incubation in complete DMEM, 0.1 μg/ml of LPS alone or with varying concentrations of VIP or VIP amphiphiles were added to the wells. After 6 hours or 24 hours of incubation, supernatants were collected for tumor necrosis factor α (TNF-α) assessment and cells were lysed by 1% Triton X-100 (Sigma) for cell population determination by PicoGreen dsDNA kit (ThermoFisher). TNF-α secretion during the experiment was evaluated by TNF-α enzyme-linked immune sorbent assay (ELISA) kit (Biolegend).

Bone marrow cells were isolated and purified from mice following an already established protocol [1]. A 10 mL solution of complete RPMI media (RPMI 1640 medium (ThermoFisher), 10% FBS (Sigma Aldrich), 1% penicillin/streptomycin (ThermoFisher)) supplemented with 20 ng/ml GM-CSF (Granulocyte-Macrophage Colony Stimulating Factor, R&D systems) was added to freshly harvested murine bone marrow cells in 100 mm diameter non-treated petri-dishes. On day 3, another 10 mL of GM-CSF-supplemented complete RPMI media was added to the previous media to avoid disturbing the cells. On day 6, cells were treated with enzyme free cell dissociation buffer (ThermoFisher) and split into two 100 mm diameter petri-dishes. Another 10 mL of GM-CSF-supplemented complete RPMI media was added to each petri-dish on day 8. On day 10, cells were harvested using enzyme free cell dissociation buffer and then counted and seeded (0.1 million per well) in new 24-well plates. After overnight incubation, 0.1 μg/ml of LPS alone or with varying concentrations of VIP or VIP amphiphiles were added to the wells. After 6 hours or 24 hours of incubation, supernatants were collected for TNF-α (mouse TNF-α ELISA, Biolegend) and CCL22 (mouse CCL22 ELISA kit, Sigma-aldrich) assessment. Cells were collected by employing enzyme free cell dissociation buffer and stained with PE - anti CD86, PE/cy7 - anti CD11c, and APC - anti MHCII (Biolegend) for surface marker detection and Live-or-Dye 488/515 Fixable Viability Staining Kit ™ (Biotium) for cell viability assessment. Cells were then analyzed by a flow cytometer (BD LSRFortessa X20) equipped with FACSDiva 8.0 Software used to detect the fluorescence intensity of the cell samples. Flowjo was utilized to gate the cells and analyze the fluorescence signal.

**Statistical analysis:** JMP software (SAS Institute) was used to make comparisons between groups using an analysis of variance (ANOVA) followed by Tukey's HSD test to determine pairwise statistically significant differences ($p < 0.05$). Within graphs, groups that possess different letters have statistically significant differences in mean whereas those that possess the same letter have statistically insignificant differences.

### Supporting Figures

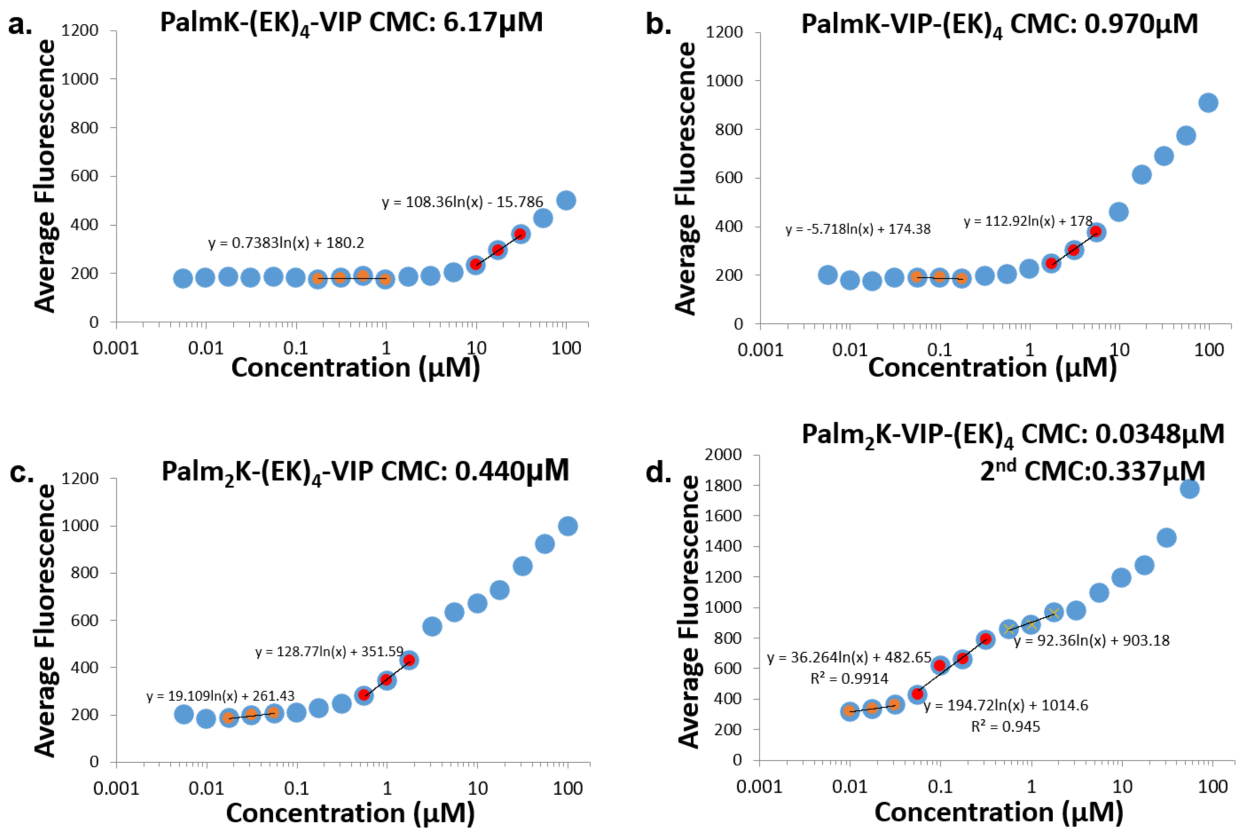


**Figure S1.** Critical micelle concentration determination for PalmK-$(EK)_4$-VIP (a), PalmK-VIP-$(KE)_4$ (b), $Palm_2K$-$(EK)_4$-VIP (c), and $Palm_2K$-VIP-$(KE)_4$ (d).

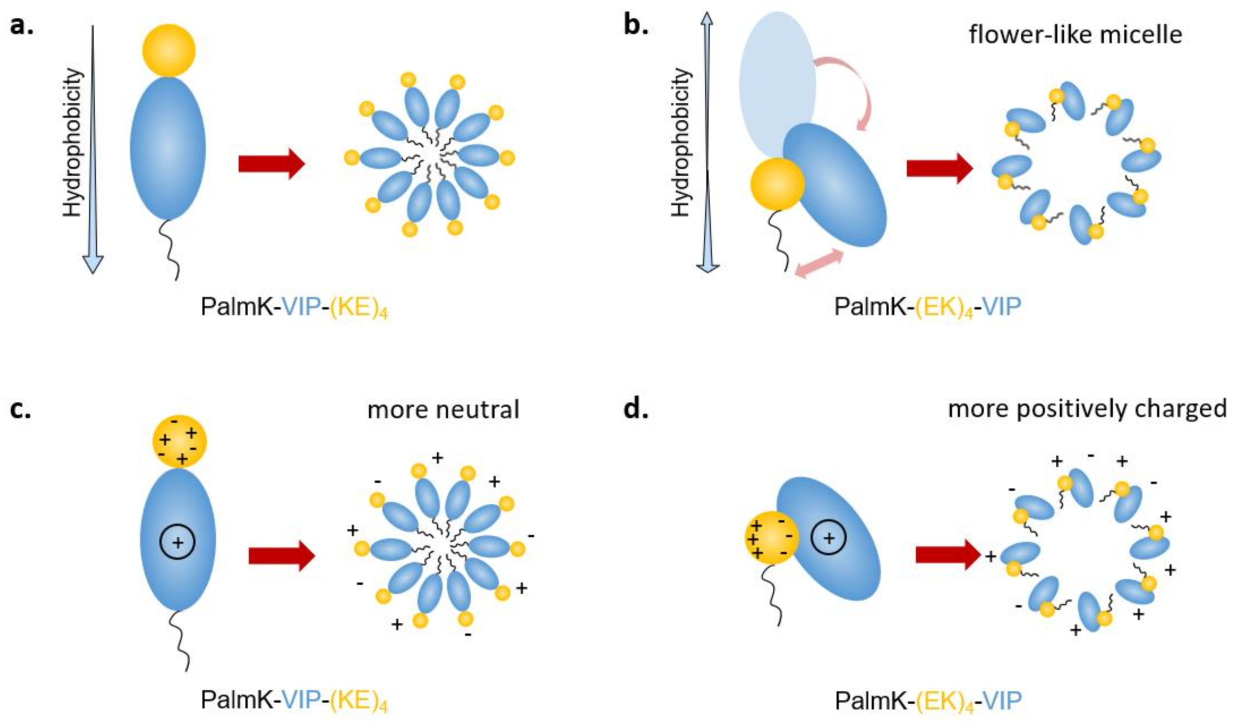


**Figure S2.** Conformation accommodation of zwitterion-like peptide external block (a & c) and internal block (b & d) peptide amphiphiles undergoing hydrophobic self-assembly (a & b) and showing charge distribution/association (c & d). The yellow ellipse, blue ellipse, and black line denote the zwitterion-like peptide block, the VIP block, and the lipid block, respectively.

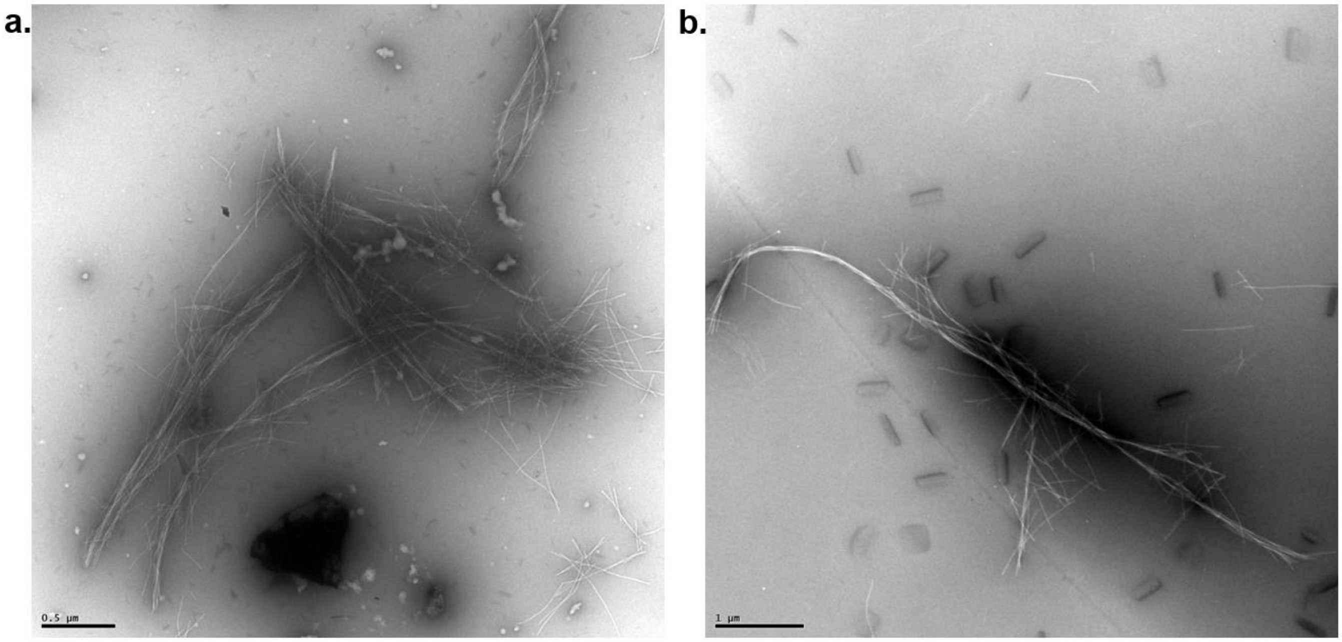


**Figure S3.** Morphology of PalmK-$(EK)_4$-VIP (a) and PalmK-VIP-$(KE)_4$ (b) at 100 µM in phosphate buffered saline (PBS) as captured by transmission electron microscopy (TEM). Micrographs were taken at magnifications of 5,000 x and 3,000 x with scale bars representing 0.5 µm and 1 µm, respectively. These images reveal the formation of long cylindrical micelles that further aggregated together into braid-like structures.

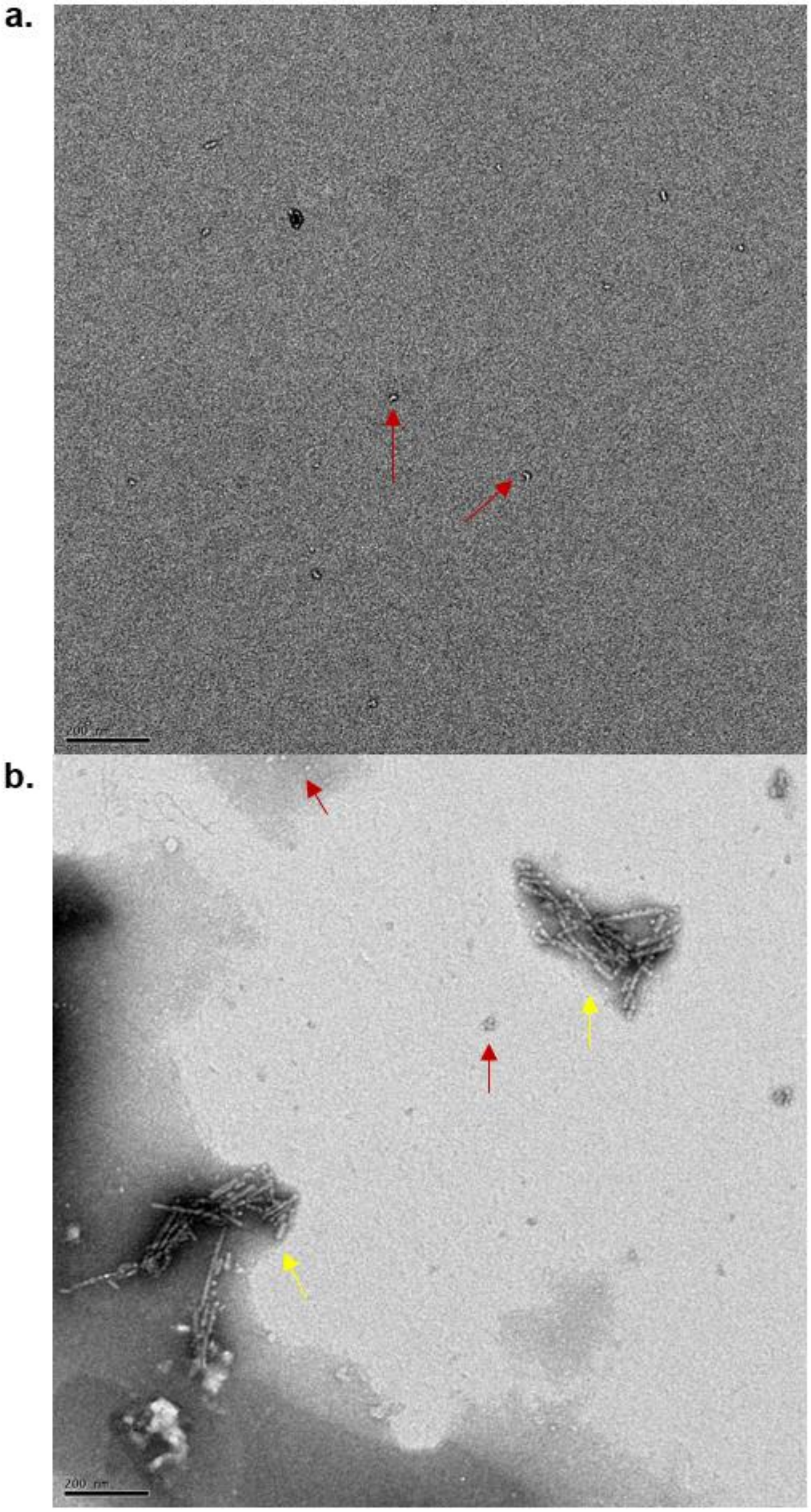


**Figure S4.** Morphology of Palm$_2$K-VIP-(KE)$_4$ structures at 0.1 μM (a) and 1 μM (b) in phosphate buffered saline (PBS) as captured by transmission electron microscopy (TEM). Micrographs were taken at a magnification of 12,000 x with a scale bar for both images representing 200 nm. Only spherical micelles were observed at 0.1 μM (red arrows) whereas both spherical micelles (red arrow) and cylindrical micelles (yellow arrow) were observed at 1 μM.

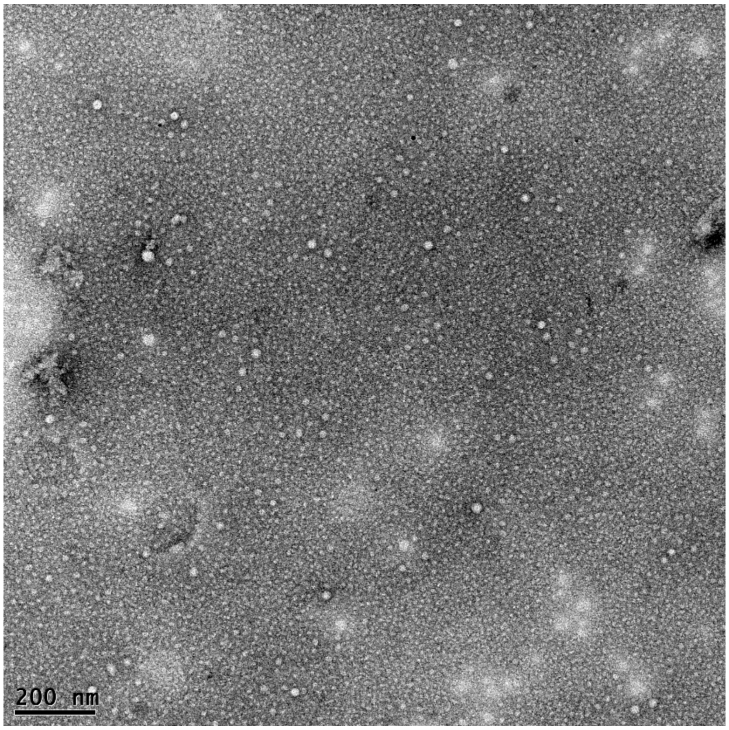


**Figure S5.** Morphology of VIP structures at 100 μM in phosphate buffered saline (PBS) as captured by transmission electron microscopy (TEM). Micrographs were taken at a magnification of 12,000 x with a scale bar for image representing 200 nm. Spherical nanoparticles were observed at this concentration.

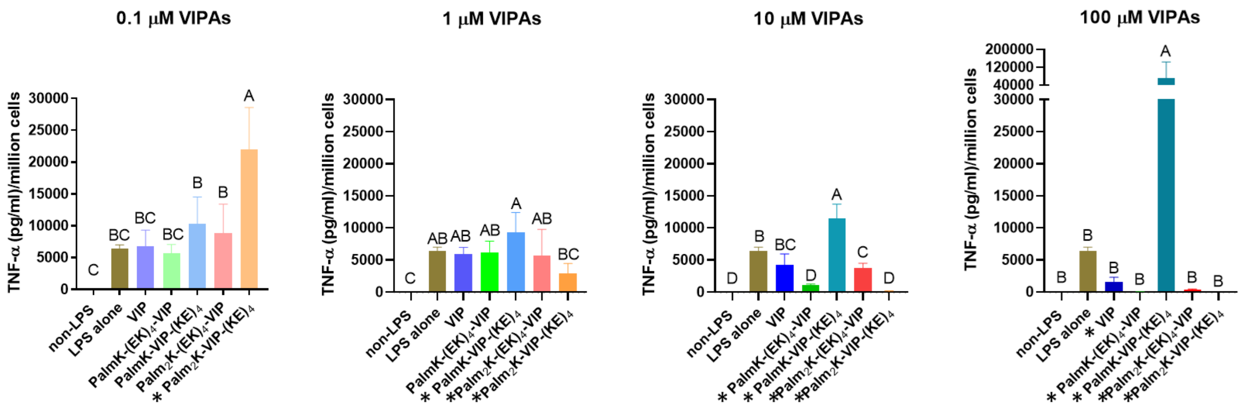


**Figure S6.** Effect of various VIPA formulation and concentration on LPS-activated Mφ TNF-α secretion after 6 hours of treatment normalized on a per million cells basis. In each graph, groups possessing the same letter have no statistically significant difference ($p > 0.05$).

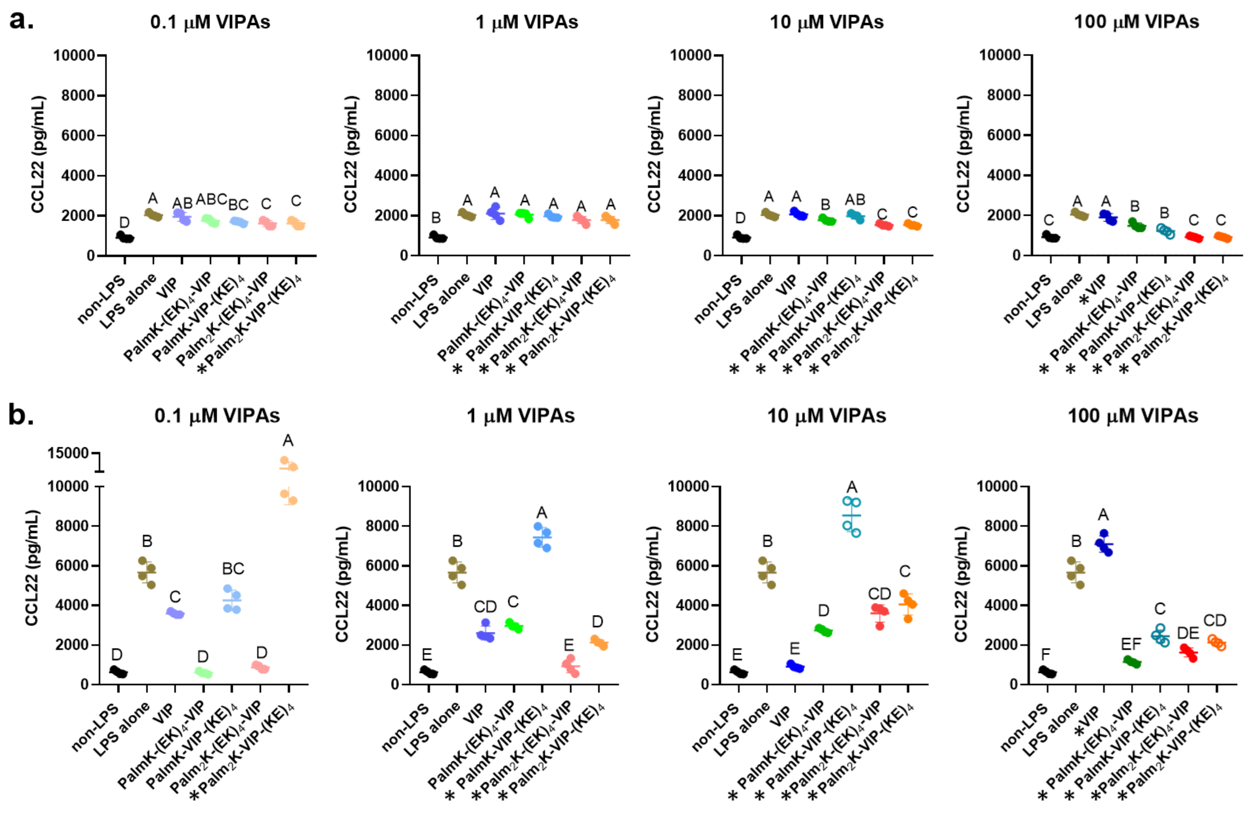


**Figure S7.** VIPAs impacts the CCL22 secretion on DCs after 6 and 24 hours of incubation. In each graph, groups possessing the same letter have no statistically significant difference ($p > 0.05$). The "*" sign in front of the name of the VIPA formulation indicates it is above the CMC and micelles have been formed.
